# MEMS-Actuated Carbon Fiber Microelectrode for Neural Recording

**DOI:** 10.1101/385153

**Authors:** Rachel S. Zoll, Craig B. Schindler, Travis L. Massey, Daniel S. Drew, Michel M. Maharbiz, Kristofer S. J. Pister

## Abstract

Microwire and microelectrode arrays used for cortical neural recording typically consist of tens to hundreds of recording sites, but often only a fraction of these sites are in close enough proximity to firing neurons to record single-unit activity. Recent work has demonstrated precise, depth-controllable mechanisms for the insertion of single neural recording electrodes, but these methods are mostly only capable of inserting electrodes which elicit adverse biological response. We present an electrostatic-based actuator capable of inserting individual carbon fiber microelectrodes which elicit minimal to no adverse biological response. The device is shown to insert a carbon fiber recording electrode into an agar brain phantom and can record an artificial neural signal in saline. This technique provides a platform generalizable to many microwire-style recording electrodes.

## I Introduction

Intracortical microelectrodes for neural recording are a powerful tool for capturing the activity of individual neurons, enabling further understanding of neural patterns that could indicate underlying disease, aid in fundamental neuroscience research, and enable brain-machine interface technologies. Current state-of-the-art electrode arrays consist of several tens to hundreds of recording sites [1]–[4]. Neural recording arrays are typically implanted to a target depth in a given region of interest. Minor adjustments are then made to the array’s depth or position to maximize the number of recorded units across the array; still, many sites may not detect active units. Devices such as the Utah array and tungsten microwire arrays are limited in that all recording sites are at a fixed relative position within the array. Upon implantation, individual recording sites cannot be independently inserted to unique depths to maximize recorded unit activity on each electrode [5], [6].

Further, within the days or weeks following implantation, the adverse biological response elicited by the implanted array may degrade the recorded signal quality [7]. Recent work has suggested that recording electrodes with cross-sections on the order of single-digit microns do not show significant evidence of neuronal loss, gliosis, or macrophage activation [2], [8]–[11]. Several carbon fiber microwire neural recording arrays have been developed with electrodes on such a scale, but none yet affords independent depth control of individual recording electrodes [3], [12]–[14].

An ideal recording platform would enable actuation of each implanted electrode to an independent depth in order to maximize signal-to-noise ratio (SNR) or until an otherwise desirable spiking unit is located [15]. During initial electrode placement, these microdrives may alternate between inserting and recording neuronal activity from the electrode, providing surgeons with real-time feedback indicating optimal placement depth [16], [17].

Recent studies have demonstrated fluidic, thermal, electrostatic, and DC microactuators capable of inserting 12 μm to 150 μm diameter electrodes with depth precision ranging from 1 μm to 25 μm [16]–[21]. Only Vitale et al.’s work demonstrates insertion of electrodes sufficiently fine to minimize the adverse biological response; however, their fluidic pumps could make insertion of large quantities of electrodes difficult [18]. In comparison with thermal actuators, the actuation method presented here does not exhibit any thermal heating [17]. Although Otchy et. al’s microdrive system was able to achieve 1 μm depth precision, each microelectrode in the implanted tetrode is 200 μm in diameter and unable to be independently placed apart from each of the other microelectrodes within the tetrode [20], [22].

We report an electrostatic-based MEMS carbon fiber actuator (Fig. 1) capable of inserting electrodes to a variable depth with 1 μm to 2 μm precision. This mechanism is capable of inserting 7.2 μm carbon fiber recording electrodes which do not elicit glial scarring [2], [8]. Although this work demonstrates insertion of single electrodes, the electrostatic actuator mechanisms could be fabricated as arrays, enabling increased channel count. The actuators are controlled via a set of voltage signals generated using an Arduino Uno.

**Fig. 1.**
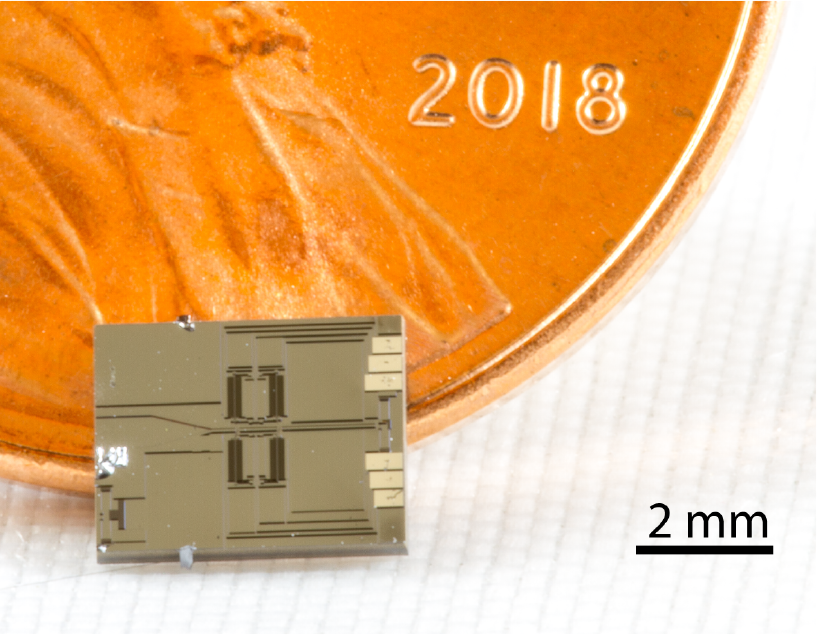
Die photo of the MEMS actuator next to a US penny. The chip is 4.5 mm x 3.5 mm, and has a mass of 22 mg.

## II Materials and Methods

### A. Theory of Operation

#### 1) Fiber Insertion

In this work, we designed a MEMS actuator containing an electrostatic motor with angled arms, as in [23], [24]. The electrostatic motors presented here are capable of producing millinewton forces over many millimeters of travel [23]–[26], sufficient for the penetration forces (calculated to be on the order of hundreds of micronewtons for this electrode style) and depths necessary for most applications [2], [3], [8]. In prior work, these actuators have been used to advance 7.2 μm carbon fibers in air, but have not been characterized for their mechanical insertion and electrical characteristics in the context of neural recording [24].

Each actuation cycle of the electrostatic motor pushes the fiber a small distance. Motion is achieved by applying voltage to an interdigitated set of capacitive fingers. Initially, one set of fingers is grounded, while the other set is held at V_*actuate*_. This electrostatic force causes the interdigitated capacitive fingers to pull in towards one another, in turn pushing out a set of flexible angled arms which grip the carbon fiber. To disengage the flexible arms from the carbon fiber, both sets of capacitive fingers are grounded and a spring pulls the capacitive fingers apart.

By using two such actuators to perform a cyclic motion in which the angled arms come into contact with the carbon fiber, move it forwards one step, disengage, and return to their initial position, the motor accumulates small steps which eventually advance the microelectrode over a large distance. For more details on actuation, see [23].

Fig. 2 shows equivalent circuit diagrams (left) and corresponding images of the device (right) with electrically connected segments labelled and highlighted.

**Fig. 2.**
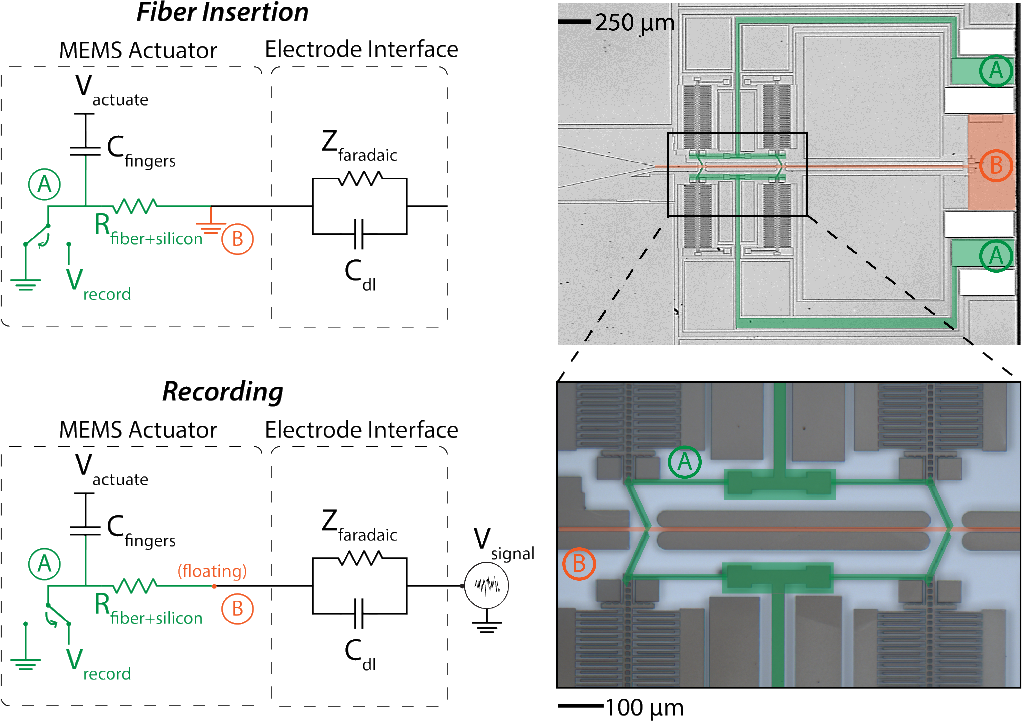
Left: Equivalent circuit models of the device. During fiber insertion (top), the electrostatic actuator pushes the fiber forwards in channel. During recording (bottom), neural signals are captured by closing all four angled arms until they contact the carbon fiber. Electrically connected segments have been labelled and highlighted. “A”, green, represents the silicon traces and angled arms which contact the carbon fiber; “B”, orange, represents the carbon fiber channel and substrate. Right: Image of the device, with inset shown at bottom.

During insertion of a fiber (Fig. 2, top left), the silicon angled arms and one set of capacitive fingers in each actuator are tied to ground (highlighted in green, “A”). The other set of capacitive fingers alternates between V_*actuate*_ and ground, dictating whether the angled arms are in contact with or disengaged from the carbon fiber. The substrate (highlighted in orange, “B”) is also grounded to prevent released silicon structures from electrostatically pulling in to the substrate.

In Fig. 2 (top right), the green highlighted regions indicate the signal path from the wirebond pads, through the silicon traces, to the angled arms which contact the carbon fiber. The orange highlighted regions indicate the substrate connection and location of a carbon fiber within the channel. Fig. 2 (bottom right) shows an inset of the silicon traces and angled arms (green) which come in contact with the fiber, nominally held in place in the fiber channel (orange).

When no voltage is applied to the actuator motor, the carbon fiber is not in contact with any silicon structure other than the substrate.

#### 2) Recording

When recording signals from the electrode (Fig. 2, bottom left), all four angled arms make contact with the carbon fiber when a high voltage, V_*actuate*_, is maintained across the capacitive fingers. These silicon arms, along with the corresponding silicon routing, form a signal path with which to record the neural signal. The path of impedance for this signal, from the tip of the carbon fiber in contact with the electrolyte to the external sensor circuity wire-bonded to the die, includes: parallel double-layer capacitance C_*dl*_ and faradaic impedance Z_*faradaic*_ at the electrode-electrolyte interface; carbon fiber impedance R_*fiber*+*silicon*_; contact resistance between the carbon fiber and silicon angled arms; silicon traces; and wire bonds (all included in R_*fiber*+*silicon*_). The electrophysiological potential is recorded from the signal pad (green, “A”) versus a reference electrode. The substrate and carbon fiber channel (orange, “B”) are left floating to prevent grounding of the recorded signal. Although the voltage difference between the sets of capacitive fingers becomes V_*actuate*_ − V_*signal*_ due to the micro-to-millivolt amplitude of neural recordings, this voltage is still sufficient to allow the angled arms to grip the fiber.

### B. Device Fabrication and Assembly

The MEMS actuator was fabricated with a two-mask silicon-on-insulator (SOI) process. Commercial SOI wafers consisting of a silicon substrate (550 μm), buried oxide layer (2 μm), and a device silicon layer (40 μm, 3250 Ω/□), were used for all devices. First, aluminium was evaporated onto the wirebond sites to improve bond adhesion. Device silicon was lithographically patterned and etched using a deep reactive ion etch (DRIE). A subsequently patterned through-etch of the silicon substrate layer, also via DRIE, served to singulate the devices. Finally, a timed vapor HF etch was used to release the structures. The fabricated device is shown in Fig. 1 and Fig. 2 (right).

Electrical connections were made by wirebonding signal wires from the chip to an off-chip set of interconnects. A substrate grounding wire and the chip were held in place with silver epoxy. Placement of the fiber within the channel was achieved by adhering a fiber to a silicone-coated tungsten micromanipulator probe tip. The probe tip and carbon fiber assembly was aligned to the left of the device layer funnel, lowered to the correct height above the chip, and inserted into the channel.

The chip is 4.5 mm by 3.5 mm, and has a mass of 22 mg. The actuator/motor area is approximately 1.5 mm^2^.

### C. Penetration into Agar

An agar brain phantom with 0.6% w/w concentration was chosen to mimic mechanical properties of the brain [27], and was used to test the penetration capabilities of carbon fibers driven by the MEMS actuator. As shown in Fig. 3, the agar was placed 400 μm from the right edge of the MEMS chip. The fiber was advanced using the minimum voltage *V_actuate_* needed to move the fiber (in the range of 20 V to 70 V, typically about 55 V). This work did not attempt to prevent discharge or shorting of high-voltage nodes with biological surfaces, but is further discussed in the conclusion of this study as future work.

**Fig. 3.**
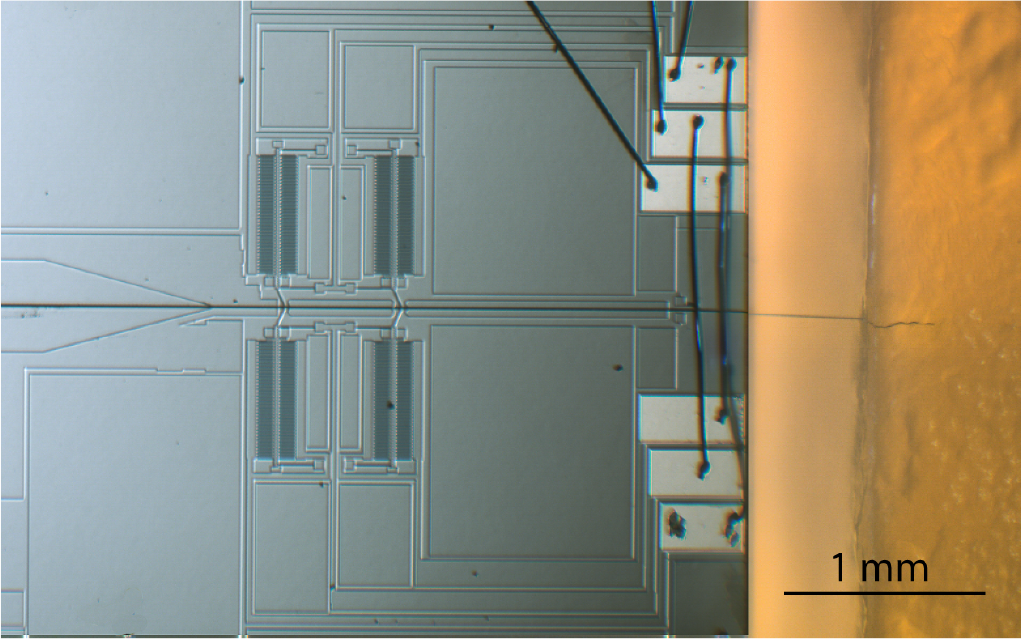
The MEMS actuator and a 7.2 μm diameter carbon fiber inserted approximately 400 μm into the agar brain phantom.

### D. Electrical Impedance Spectroscopy

Electrode impedance measurements were conducted using a Keysight E4980AL precision LCR meter. Measurements were obtained by applying a 1V_*rms*_ signal of varying frequency between the MEMS signal recording pad and a tungsten electrode placed in 10X PBS. The carbon fiber which extended beyond the MEMS chip edge was also placed in the 10X PBS solution. To ensure proper electrical contact with the fiber, as would be the case during a neural recording, angled arms were kept in contact with the carbon fiber by applying a high voltage to the interdigitated fingers. Frequency was swept over 21 increments from 100Hz to 10KHz, with three measurements taken at each frequency increment and subsequently averaged.

### E. Playback of Neural Recording

To simulate an *in vivo* recording, a dataset previously recorded from a microwire in an awake/behaving rat motor cortex was played back over a waveform generator (Analog Discovery 2) onto a platinum “neural signal” electrode in 1X PBS. A silver reference electrode and the tip of a carbon fiber (the “recording electrode”) held by the microelectrode actuator were also placed in the PBS to form a complete circuit. Signals recorded by the microelectrode actuator were amplified using a DAM50 bio-amplifier (World Precision Instruments) and digitized using an Agilent Technologies DSO-X 3034A digital oscilloscope.

## III Results and Discussion

### A. Penetration into Agar

With *V_actuate_* = 55 V applied across the interdigitated fingers at a frequency of 20 Hz, the carbon fiber was successfully able to penetrate 400 μm into an agar brain phantom (Fig. 3). Since the fiber travelled 400 μm from the edge of the chip to the agar, the distance travelled was 400 μm in air and 400 μm in agar, for a total distance travelled of 800 μm. The motor was able to advance the fiber in 1 μm increments. This depth precision is dependent on the angled arm geometry and distance between opposing sets of the capacitive fingers, which in turn are limited by the photolithographic tools.

Theoretically, the actuator presented here can output over 1 mN of force at 85 V, although the experimental force output was not directly measured in this study. A previous iteration of this actuator was capable of advancing a carbon fiber up to 1.8 mm in air [24]. A higher force-output version of the actuator presented here should be able to advance a carbon fiber a similar distance into agar. At this voltage and frequency, the actuator consumes tens of microwatts of power [26].

The carbon fiber is supported on three sides: by the substrate from underneath, and on both sides by the two 40 μm silicon sidewalls which form the fiber channel. Once the fiber advances beyond the edge of the chip and dimples the agar, it can be considered as a fixed-guided beam, as the fiber is no longer free to move laterally at the contact point [8], [12].

Fig. 4 shows a carbon fiber inserting into the agar brain phantom. These snapshots of an insertion event suggest that, when operated at 20 Hz, the motor is capable of inserting the electrode at rates up to 10.5 μm*/*s. This style of electrostatic motor has been shown to operate at speeds of up to 30 mm*/*s in air when operated at 8 kHz [26], providing an upper bound on the theoretical insertion speed.

**Fig. 4.**
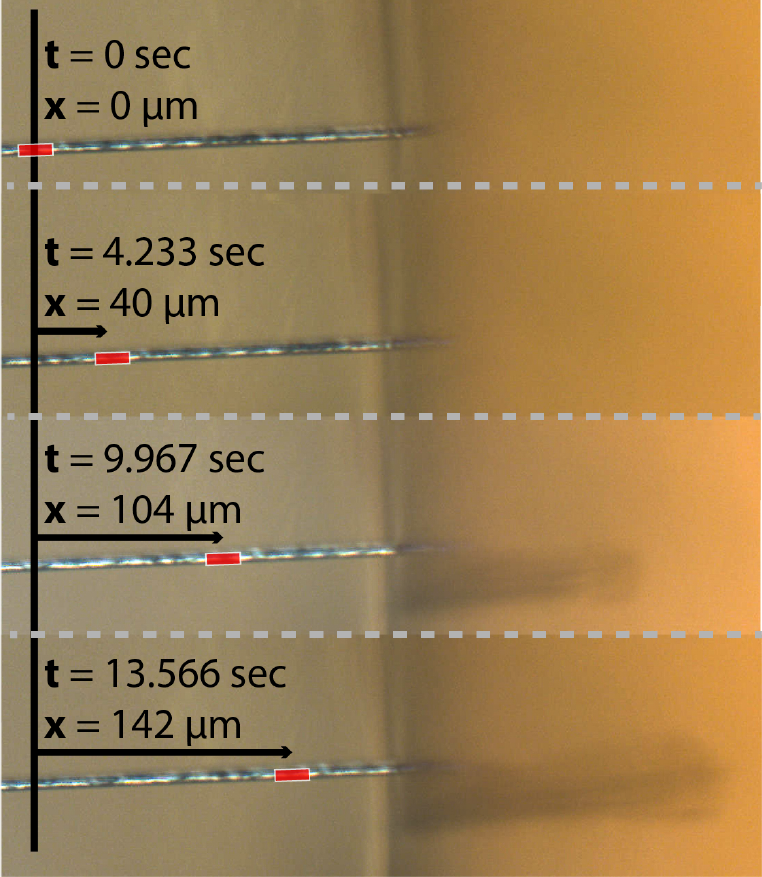
Multiple close-up shots showing the insertion of a 7.2 μm diameter carbon fiber into the brain phantom. The red dash indicates the same position on the carbon fiber between frames.

Motor speed exhibits a linear dependence on the operating frequency of the high-voltage waveforms applied to the electrostatic fingers [26]. Slippage was observed at the interface between the carbon fiber and the silicon angled arms, although the degree to which slippage occurred was not quantified. In future work, slippage can be minimized by increasing the force output of the motors either by changing the motor design or operating the electrostatic fingers at a higher voltage. Additionally, modifying the angled arm contact surface geometry or insulating portions of the fiber to increase static friction with the carbon fiber may aid in decreasing slippage.

As a side effect of working in an open, non-cleanroom workspace, small dust particles often became stuck in the channel, necessitating short bursts of repeated voltage ramping to force the fiber past dust particles. From a practical perspective, build-up of dust particles in the channel is the only aspect of this setup which prevents a single actuator from being used to advance longer or multiple electrodes.

### B. Electrical Impedance Spectroscopy

Fig. 5 shows the impedance spectroscopy for a single carbon fiber electrode held by two angled arms in the actuator fiber channel. Note that the magnitude of the impedance halves when four angled arms are in contact with the fiber, rather than two. The constituent components of this lumped electrode impedance include the electrolyte-electrode double-layer impedance C_*dl*_ and Z_*faradaic*_, impedance of the 10 mm-long carbon fiber R_*fiber*_, contact resistance between the carbon fiber and the silicon angled arms, resistance of the silicon traces R_*silicon*_, and resistance of the wirebonds and external wires which lead to an off-chip ADC. The resistance of the silicon traces dominated upon measuring each of the constituent impedances in isolation. Future work includes metallizing the silicon traces to reduce sheet resistance from 3250 Ω/□ with bare silicon traces to approximately 0.3 Ω/□ for 100 nm aluminum-coated traces, helping to decrease the total silicon trace resistance to approximately 1 kΩ. The total capacitance of *C_fingers_* was not directly measured, but is theoretically calculated to be 7.5 pF.

The double layer impedance between 1X PBS and a 5 mm-long, 7.2 μm-diameter fiber was approximately 15 kΩ, and at low frequencies was mostly resistive in phase. At higher frequencies, the magnitude of impedance dropped to 5 kΩ, which matches the 1 kΩ*/*mm to 2 kΩ*/*mm expected impedance of the fiber alone. This result is a function of the electrode area exposed to electrolyte; in this case, 0.11 mm^2^. This is thousands of times larger than in a typical recording scenario in which only the tip of the carbon fiber is exposed to the electrolyte. Additionally, in a typical recording scenario the recording site would be electroplated to decrease its impedance by up to two orders of magnitude [8], [12].

**Fig. 5.**
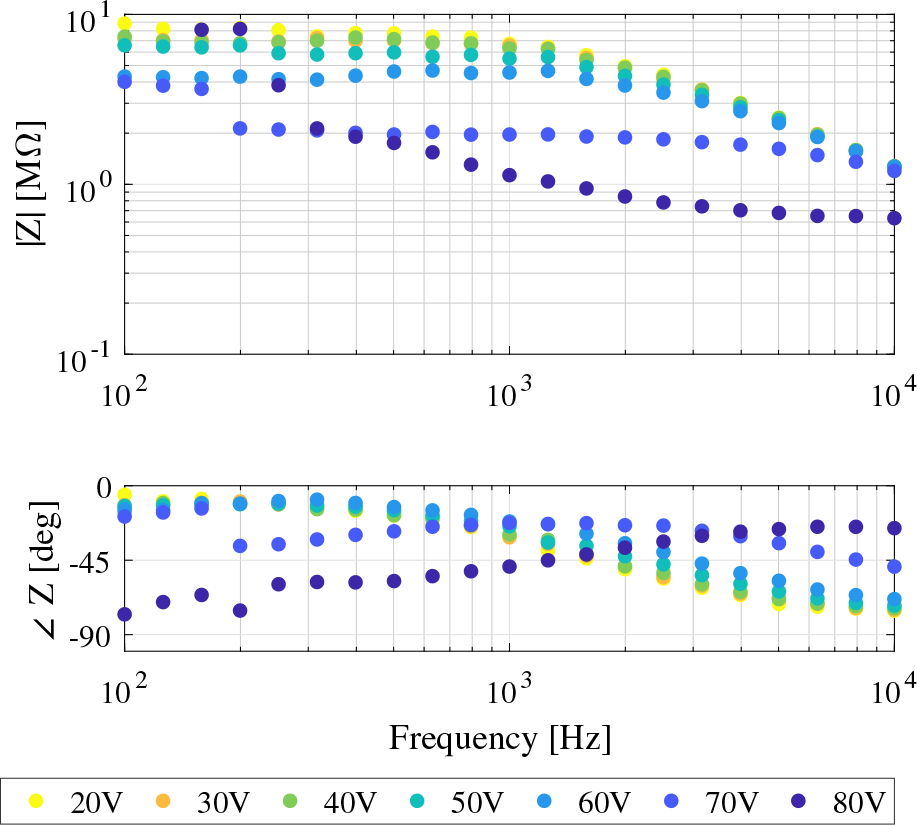
Electrical impedance spectroscopy for a single electrode. The microelectrode is held in place by two angled arms. Upper plot shows impedance magnitude; lower plot shows impedance phase. Different colors represent different actuation voltages applied to the electrostatic fingers. Impedance is dominated by the resistance of the silicon traces.

Each of the silicon traces leading from the angled arms to the wirebond pads leading off chip was 10 MΩ. With all four traces in parallel, the resistance drops to 2.5 MΩ. The contact resistance between a single silicon pawl and the carbon fiber is on the order of 100 kΩ; with four angled arms contacting the carbon fiber, the resistance decreases to 25 kΩ. Overall, the electrode-electrolyte, carbon fiber, and carbon fiber-silicon contact impedances are negligible as compared to that of the silicon traces, but in a scenario in which the traces are metallized and the recording site is small, the electrode-electrolyte impedance should dominate.

Fig. 5 also demonstrates significant capacitive crosstalk above 10 kHz. This is likely due to the the silicon trace resistance, which can be decreased by a factor of 10,000 by metallizing the traces, pushing the crosstalk effect out to significantly higher frequencies.

As observed in the magnitude plot of Fig. 5, the impedance magnitude decreases as the actuation voltage increases. This is likely because the angled arms grip the fiber with greater force at greater actuation voltages, reducing the contact resistance between the angled arms and carbon fiber electrode.

### C. Playback of Neural Recording

To mimic *in vivo* recordings from a mouse, multi-unit activity previously recorded from a microwire in an awake/behaving rat motor cortex was played back and recorded using the microelectrode actuator in “record” mode. As seen in Fig. 6, the carbon fiber recording closely resembles that of the signal played back by the waveform generator. The maximum signal amplitude is 70 mV. Although some minor features seen in the waveform generator signal are lost in the carbon fiber recording, major spikes putatively corresponding to firing of the mouse neurons are still clearly visible.

**Fig. 6.**
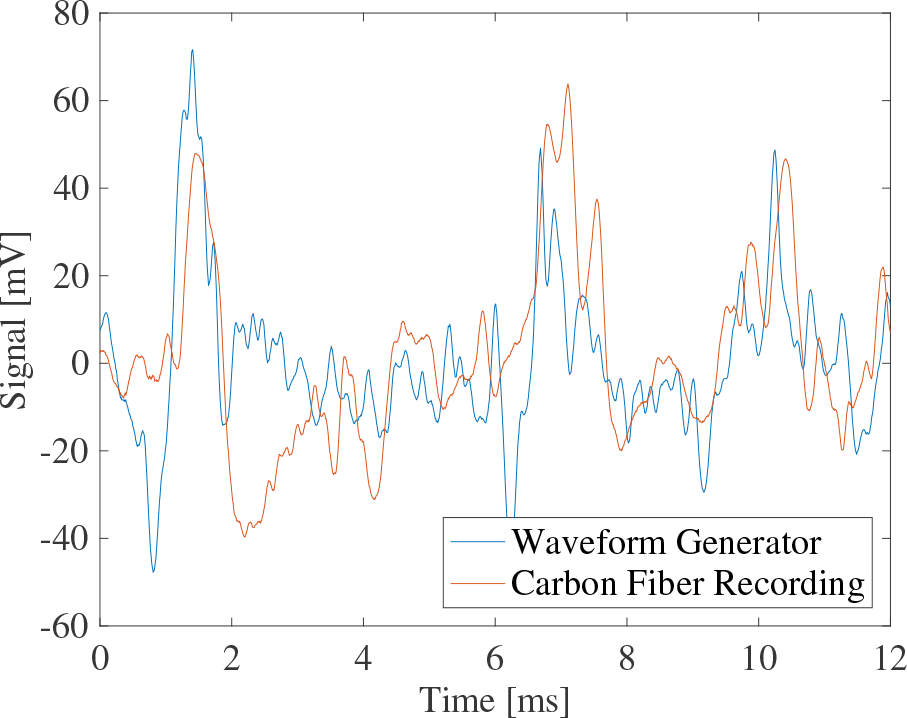
Playback of multi-unit activity (blue) is recorded by the carbon fiber microelectrode (red) in 1X PBS. The data was sampled at 125 kHz and a ten sample moving average filter was used to smooth the data.

## IV Conclusions and Future Work

Electrostatic MEMS microelectrode actuators offer a platform to insert carbon fiber filaments into neural tissue with micron-precision. Based on our measured data, the current actuator is capable of inserting fibers up to 400 μm into an agarose brain phantom at estimated speeds of up to 10.5 μm/s. By taking advantage of previously-designed electrostatic gap closer mechanisms [23], [24], [26], this design could be developed to achieve force outputs necessary to push fibers even greater distances, with greater step precision and greater speed.

The electrical characteristics of the actuator mechanism in the “record” mode are suitable for neural recording applications. Although the current design’s impedance is dominated by the silicon traces for a net impedance of 4 MΩ to 8 MΩ, this overall impedance can be decreased by metallizing the silicon traces. While this study used uninsulated carbon fiber filaments, electrodes viable for recording would be insulated in parylene-C near the recording site. Additionally, electroplating poly(3,4-ethylenedioxythiophene) doped with polystyrene sulfonate (PEDOT:PSS) on the recording site would significantly reduce electrode impedance and improve recording characteristics [3], [8].

Layout area could further be minimized to allow minimum-pitch arrays of electrostatic actuator mechanisms to simultaneously position multiple 7.2 μm carbon fibers. Further, creating a bidirectional actuation mechanism would require either duplicating the existing motor or creating a mechanism to reverse the direction of angled arms.

Coating the actuator’s capacitive fingers with a non-conductive material such as alumina would prevent the high-voltage and grounded fingers from accidentally shorting together and discharging through the carbon fiber [28]. Additional packaging is necessary to isolate the electrically active wirebond pads.

This device could also be used as a general platform for flexible electrode insertion. Recently, Luan et al. used a 7 μm carbon fiber to mechanically support 10.5 μm by 1.5 μm flexible recording electrodes during insertion [29]. The actuators presented in this work could be placed on the surface of the brain and used to inject the electrodes developed by Luan et al. [29]. With enlargement of the fiber channel, this device could also be used to deliver optical fibers to precise depths in the brain for optogenetics studies [30].

## Acknowledgment

We thank the Berkeley Sensor & Actuator Center and the Chan Zuckerberg Biohub for support during this project. All fabrication was performed at the UC Berkeley Marvell Nanolab. The authors would also like to thank D. Contreras, J. Edmunds, H. Gomez, J. Greenspun, R. Muller, D. Piech, K. Shen, R. Shih, T. Zajdel, and A. Zhou for their support throughout this project.

